# Asymmetry and redundancy of STAT5 paralogs across CD8^+^ T cell differentiation states

**DOI:** 10.1101/2025.07.23.666302

**Authors:** Svetlana Ristin, Molly Dalzell, Christopher Armstrong, Nisa Ilsin, Antonio M. Fontanella, Luis Nivelo, Lothar Hennighausen, John J. O’Shea, Alejandro V. Villarino

## Abstract

Fostering STAT5 signaling is key to immunotherapies that leverage CD8^+^ T cell biology. Using mouse models, we demonstrate that the two mammalian STAT5 paralogs, STAT5A and STAT5B, are at once redundant and functionally distinct in CD8^+^ T cells. Specifically, we establish that they are *asymmetric paralogs*, exhibiting both widespread homology at molecular level and functional asymmetry at cellular level, with STAT5B emerging as dominant. In fact, compared to STAT5A, STAT5B deficiency had greater impact on nearly all parameters tested. As a mechanism, we determined STAT5B is twice as abundant, accounting for two-thirds of the total STAT5 pool. We also defined both cytokine- and cell state-restricted STAT5B functions, and a core gene signature that highlights universal effects. Together, these studies affirm the centrality of STAT5 in CD8^+^ T cells, reveal common and circumscribed activities, and present a unifying model for paralog redundancy that foregrounds and explains the dominance of STAT5B.

**Summary**: STAT5 paralog dominance and redundancy in CD8^+^ T cells

## Introduction

STAT5 is a signal-dependent transcription factor that mediates key cellular pathways in immune cells, ranging from universal, pan-lineage processes like glucose metabolism and apoptosis, to specialized, lineage-specific processes like cytokine production and cytotoxicity (1–6). STAT5 has been most extensively studied in lymphocytes, especially T cells, where it exerts numerous pro- and anti-inflammatory functions. Examples of the former include its ability to promote CD8^+^ and NK cell cytotoxicity (7–10), and its ability to promote IFN-*γ* production across the lymphoid compartment (2, 3, 10–12). Examples of the latter include its ability to marshal CD4^+^ regulatory T cells (Treg) and to suppress Th17- and follicular helper-type (Tfh) T cell responses (13–21). Genetic studies illustrate both the importance of STAT5 and its dual pro- and anti-inflammatory nature. Complete germline ablation in mice results in severe lymphopenia and anemia leading to perinatal death, and the few animals that survive beyond one month eventually develop lethal, systematic autoimmunity (22–25). Partial germline STAT5 deficiency also manifests a paradoxical combination or lymphopenia and autoimmunity, only less severe and, in mice, pathology is mainly localized to kidneys (26–30).

STAT5 is unique among STAT family members in that it is a collective of two proteins, STAT5A and STAT5B. Thus, the mammalian genome has 4 total *STAT5* alleles encoded by two adjacent loci, *STAT5A* and *STAT5B*, resulting from an evolutionarily recent duplication event (31, 32). Given their recent divergence, STAT5A and STAT5B are ∼95% homologous at protein level, which has led to persistent questions about whether they are redundant or functionally distinct. Arguments are compelling on both sides. First, genome-wide distribution studies have established that they mostly co-localize and, in turn, regulate many of the same genes (7, 33, 34). Second, in a qualitative sense, STAT5A and STAT5B deficiencies impact the lymphoid compartment in similar ways, which suggests analogous functions. For instance, both result in diminished NK, Treg and CD8^+^ T cells (7, 30, 33, 35). However, they are not similar in a quantitative sense. These and other phenotypes are more pronounced in mice lacking *Stat5b* than those lacking *Stat5a*, and most dramatic in mice lacking both *Stat5a* and *Stat5b*, which suggests cooperation and/or STAT5B-specific functions. For instance, *Stat5b*-deficient mice have far greater reductions in steady state lymphocytes, particularly Innate Lymphoid Cells (ILCs), and far greater defects in cytokine-driven T cell responses (7, 22, 30, 33, 35–37).

In humans, germline *STAT5B* mutations result in severe immunological dysfunction, including sharply reduced and/or defective NK, Treg and CD8^+^ T cell responses while, tellingly, no analogous *STAT5A* mutations have been reported (26–29). However, it bears noting that *Stat5a* deficiency is more impactful than *Stat5b* deficiency in certain cell types, including mammary epithelium (38–41), hematopoietic stem cells (42) and B cells (43) – and that both STAT5A and STAT5B are required for optimal elaboration of downstream transcriptional programs (7). Both paralogs are also necessary for optimal tetramerization, a process that further promotes STAT5 activity (7, 44, 45).

STAT5 is activated downstream of many cytokines and growth factors but, in lymphocytes, it is most associated with cytokines operating through the common *γ* chain (*γ*c) receptor and its associated Janus kinase, JAK3 (2, 46). Each member of this family, which includes IL-2, IL-4, IL-7, IL-9, IL-15 and IL-21, employs a dedicated co-receptor subunit that pairs with *γ*c to enable both ligand-specificity and a shared ability to activate STAT5. Downstream of *γ*c and JAK3, STAT5 is required for development and homeostasis of multiple lymphoid lineages, most notably T and NK cells (5, 25, 47), and directly instructs key genes involved in lymphocyte differentiation, including *IgK* in B cells (48, 49), *RUNX3*, a transcription factor necessary for bifurcation of the CD4^+^ and CD8^+^ T cell lineages (47), and FOXP3, the ‘master’ transcription factor of CD4^+^ regulatory T cells (Treg)(15, 20, 21). Another important feature is its ability to drive terminal differentiation, the process by which lymphocytes make irreversible fate decisions which enable effector functions. That capacity is clearly evident in CD8^+^ T cells, where STAT5 promotes cytotoxicity and memory formation/maintenance while subverting ‘exhaustion’, a dysfunctional state which curtails the effector program (8, 9).

The central role of STAT5 in lymphocyte biology has motivated research on upstream cytokines as therapeutic agents. In fact, recombinant IL-2 was the first immuno-therapy (50) and numerous trials are ongoing for this and other STAT5-activating cytokines in settings of cancer and autoimmune disease, including recent efforts to: (1) engineer upstream cytokines for ‘better’ or lineage-selective STAT5 activities, (2) combine upstream cytokines with ‘checkpoint’ inhibitors, and (3) incorporate STAT5 signaling payloads in Chimeric Antigen Receptor T cell constructs (51–55). Using a genetic model, we addressed the evergreen question of whether STAT5A and STAT5B are redundant or functionally distinct, focusing on CD8^+^ T cells, principal mediators of many (if not most) immunotherapies. We conclude that they are *asymmetric paralogs*; exhibiting both functional homology at molecular level and asymmetry at cellular level. We also identify cytokine-, lineage-, and state-restricted STAT5 functions, and a core STAT5 signature that is universally evident and contains emblematic STAT5-regulated pathways, such as carbon metabolism and apoptosis. Together, these findings affirm the centrality of STAT5 in CD8^+^ T cells, uncover context-specific STAT5 activities and establish a unifying model for functional disparity between STAT5 paralogs.

## Results

### STAT5 depletion markedly impacts the CD8^+^ T cell compartment

To compare and contrast STAT5 paralogs, we generated a series of germline knockout (KO) mice with diminishing STAT5 alleles, starting with four and dwindling to one (Fig 1A)(30, 35, 39). Hereafter, we refer to each strain according to the alleles that were deleted, save for 4 allele mice, which are ‘wild type’ (*Stat5a^+/+^ Stat5b^+/+^*). By this nomenclature, AAB mice retain one allele of STAT5B (*Stat5a^−/−^ Stat5b^+/−^*), while BBA mice retain one allele of STAT5A (*Stat5a^+/−^ Stat5b^−/−^*). We previously used this approach to study CD4^+^ helper T cells and found that STAT5 deficiency strongly impacted both effector and regulatory functions, with STAT5B emerging as dominant (30). Importantly, we used the term ‘dominant’ (as we do now) to indicate that STAT5B has greater relative impact, not as used in genetics, where it refers to a ‘dominant’ allele that overrides effects of a ‘recessive’ allele. During the course of those studies, we also noted that unlike CD4^+^ T cells, the CD8^+^ T cell compartment was greatly diminished in spleens, lymph nodes and bone marrow (30), resulting in skewed CD4/CD8 ratios (Fig 1B-E). This phenotype was most pronounced in strains bearing only one STAT5 allele (AAB or BBA), suggesting a gene dose effect, and strains lacking STAT5B. Thus, CD8^+^ T cells appear acutely sensitive to reductions in STAT5 availability and more dependent on STAT5B than STAT5A.

**Fig. 1.**
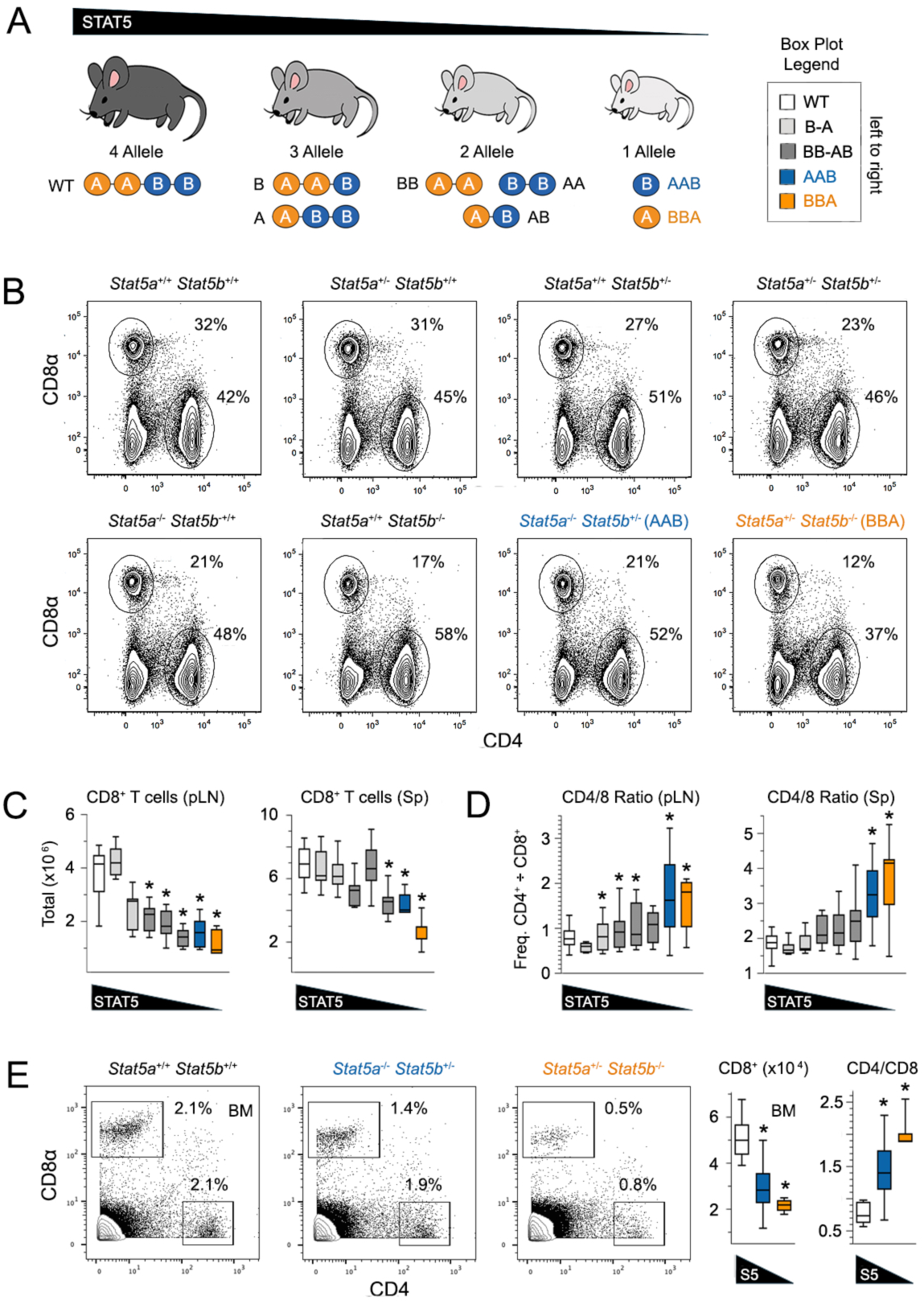
STAT5 deficiency markedly impacts the CD8^+^ T cell compartment. (A) Cartoon depicts mouse models used in this study. The 8 genotypes are grouped based on total STAT5 alleles, ranging from 4 to 1, and named according to alleles that are deleted. AAB (*Stat5a^−/−^ Stat5b^+/−^*) and BBA (*Stat5a^+/−^ Stat5b^−/−^*) mice are always colored blue and orange, respectively. (B) Flow cytometry contour plots show surface CD4 and CD8⍺ proteins in pLN. (C-D) Box plots compile cytometry-based (C) CD8^+^ T cell counts and (D) CD4/CD8 ratios from pLN and spleens. (E) Flow cytometry contour plots show surface CD4 and CD8⍺ on lymphocytes from bone marrow. Box plots compile CD8^+^ T cell counts and CD4/CD8 ratios. Replicate counts and statistical tests for all experiments are detailed in Table S6.

To determine if the observed phenotypes are T cell intrinsic, we generated mixed chimeras using bone marrow from WT and AAB or BBA donors. A 1:5 ratio of WT to KO donors was used because we expected that WT donors would have a strong competitive advantage. Indeed, despite this consideration, splenic CD8^+^ T cell ratios strongly skewed towards WT donors in the WT/AAB mix and even more so in the WT/BBA mix (Fig. 2A). By contrast, the initial 1:5 ratio was largely preserved for CD4^+^ T cells (Fig. 2B-C). In complementary studies, we also analyzed mice lacking STAT5A and STAT5B selectively in T cells and found that, along with steep reductions in total T cell counts (Fig. S1A-B), CD8^+^ T cells were conspicuously depleted in primary lymphoid organs, resulting in sharply skewed CD4/CD8 ratios (Fig. 2D). Together, these data affirm that that CD8^+^ T cells are more impacted by STAT5 deficiency than CD4^+^ T cells, that STAT5B deficiency is more deleterious than STAT5A deficiency and, importantly, that the CD4/CD8 skewing seen in our STAT5 ‘allele’ mice likely reflets T cell intrinsic functions.

**Fig. 2.**
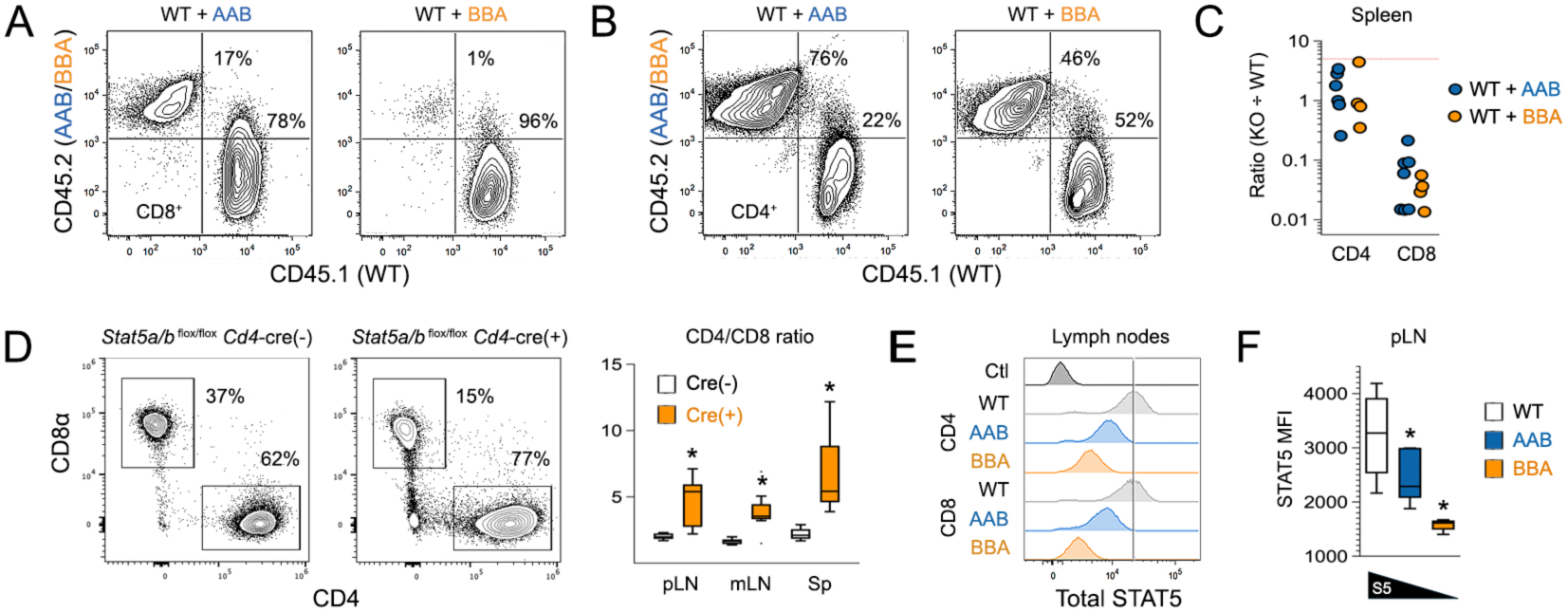
T cell intrinsic requirement for STAT5. (A-B) Flow cytometry contour plots show representative (A) CD4^+^ and (B) CD8^+^ T cell engraftment from spleens of mixed bone-marrow chimeras. WT donor cells are CD45.1 ^pos^ CD45.2 ^neg^; KO donor cells are CD45.1 ^neg^ CD45.2 ^pos^. (C) Scatter plot compiles cytometry-based engraftment data. Each point represents a discrete donor/host pair. Y axis indicates the observed KO to WT ratio in spleens. Red line denotes 5:1 KO to WT starting ratio. (D) Flow cytometry contour plots show surface CD4 and CD8⍺ proteins in pLN of *Stat5 ^flox/flox^ Cd4*-Cre^+/−^ mice and *Cd4*-Cre^−/−^ littermate controls. Box plot compiles CD4/CD8 ratios across experiments. (E) Flow cytometry histogram shows total STAT5 protein levels in splenic CD4^+^ and CD8^+^ cells of the indicated genotypes. (F) Box plot compiles total STAT5 data. Replicate counts and statistical tests for all experiments are detailed in Table S6.

STAT5B accounts for two-thirds of total STAT5 in CD4^+^ T cells (30), so we next asked if the same is true of CD8^+^ T cells. These studies showed that, indeed, far less STAT5 is detected in absence of STAT5B than STAT5A (Fig. 2E-F), leading us to conclude that relative abundance likely explains phenotypic differences between STAT5A and STAT5B deficient CD8^+^ T cells. However, relative abundance likely does not explain why CD8^+^ T cells are more sensitive to STAT5 deficiency than CD4^+^ T cells; total STAT5 content was similar between WT CD4^+^ and CD8^+^ T cells (Fig. S1C).

### STAT5 paralog asymmetry in CD8^+^ T cells

Given sharp reductions in CD8^+^ T cell counts, we next assessed the impact of STAT5 deficiency on differentiation state. Using well described surface markers, we determined that STAT5 deficiency leads to prominent accumulation of effector and memory state CD8^+^ T cells, at the expense of naive state CD8^+^ T cells. This phenotype was most pronounced in mice lacking STAT5B, and also evident mice lacking STAT5 selectively in T cells, again affirming that it is cell intrinsic (Fig. 3A-C, Fig. S2A-C). Therefore, STAT5 signaling not only enables accumulation of CD8^+^ T cells in lymphoid organs, but also helps keep them in a quiescent, naive state. STAT5A and STAT5B both participate but asymmetrically, with STAT5B emerging as dominant.

**Fig. 3.**
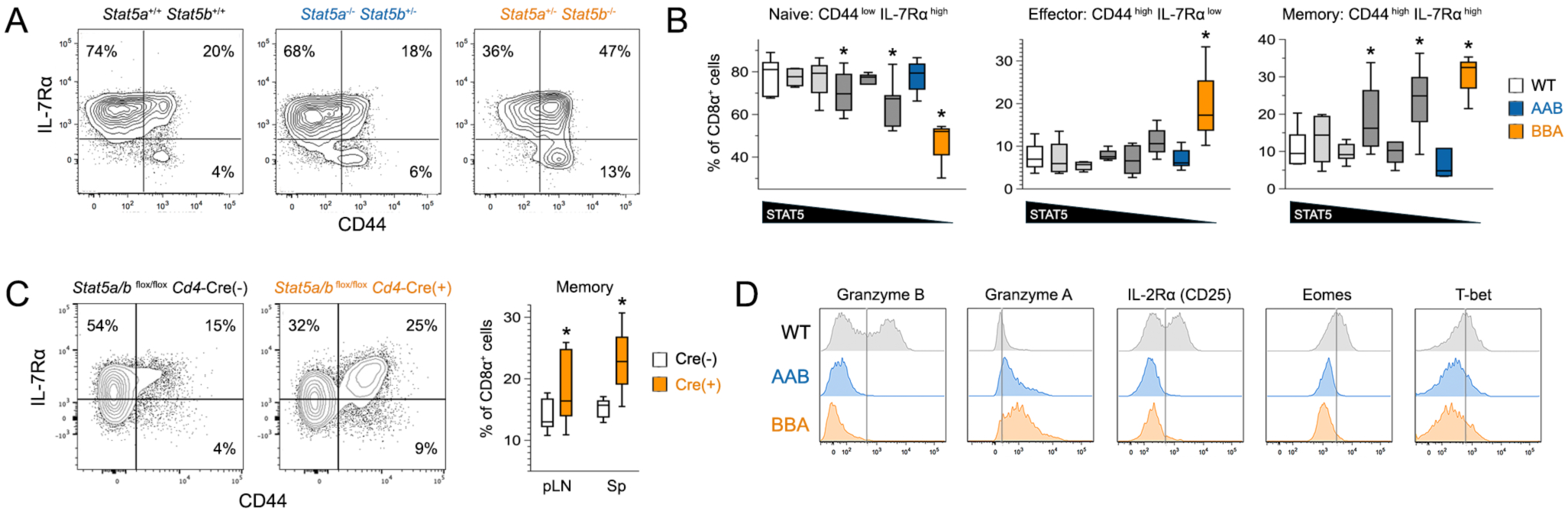
STAT5 deficiency unleashes effector and memory CD8^+^ T cells. (A) Flow cytometry contour plots show surface CD44 and IL-7R⍺ (CD127) on CD8^+^ T cells from pLN. Naïve cells are defined as CD44 ^low^ IL-7R⍺ ^high^ (upper left), central memory cells as CD44 ^high^ IL-7R⍺ ^high^ (upper right) and effector cells as CD44 ^high^ IL-7R⍺ ^low^ (lower right). (B) Box plots compile frequencies of naïve, memory and effector cells in pLN. (C) Flow cytometry contour plots show CD44 and IL-7R⍺ on CD8^+^ T cells in pLN of *Stat5 ^flox/flox^ Cd4*-Cre^+/−^ mice and *Cd4*-Cre^−/−^ littermate controls. Box plots compile frequencies of memory cells in pLN and spleens. (D). Flow cytometry histograms show surface or intracellular levels of the indicated proteins in splenic CD8^+^ T cells after 48 hours *ex vivo* activation and culture. Ex vivo culture conditions detailed in Table S7. Replicate counts and statistical tests for all experiments are detailed in Table S6.

STAT5 controls expression of key elements in the CD8^+^ T cell effector program, including granzymes, cytokine receptors and transcription factors, such as T-box family members Eomes and T-bet (7, 33). To determine if these are compromised in the absence of STAT5A and/or STAT5B, we purified naive CD8^+^ T cells from WT, AAB or BBA mice, activated them with agonist anti-T Cell Receptor (TCR) and anti-CD28 antibodies in the presence of IL-2, a potent STAT5 stimulus, then measured effector proteins by flow cytometry. Strikingly, we found that Granzyme B, IL-2R⍺, Eomes and Tbet were all sharply reduced when STAT5A or STAT5B was deleted, while Granzyme A was increased (Fig. 3D). Thus, while prone to effector differentiation, STAT5-deficient CD8^+^ T cells cannot elaborate key elements of the cytotoxic program and, in this capacity, STAT5A and STAT5B appear similarly important.

To further assess the impact of STAT5A and STAT5B deficiencies, we compared transcriptomes directly *ex vivo*, or upon stimulation with IL-7 or IL-15 (Fig. 4A, Fig. S3A). First, using standard surface markers, we sorted naïve (CD44 ^low^ CD62L ^high^), effector (CD44 ^high^ CD62L^low^) and memory (CD44 ^low^ CD62L ^high^) CD8^+^ T cells from pooled lymph nodes and spleens of AAB and BBA mice. Then, we called Differentially Expressed Genes (DEG) relative to WT controls and found striking disparity; STAT5B deficient cells (BBA) had many more DEG than STAT5A deficient cells (AAB), regardless of state or stimulus (Fig. 4B, Table S1-S2). Importantly, DEG called only in STAT5B deficient cells included many emblematic STAT5 targets (e.g. Bcl2, MYC; Fig. 4B) linked to cellular pathways known to be STAT5 driven, including several relating to T cell memory and metabolism (Fig. 4C, Fig. S3B). We also noted that IL-15 mobilized more DEG than IL-7 regardless of genotype or differentiation state, in line with a recent survey of c*γ* cytokines in immune cells (56), and that it mobilized more DEG in memory than naïve CD8^+^ T cells (Fig. 4B). Thus, STAT5B does the heavy lifting downstream of both IL-7 and IL-15, and in both naïve and memory cells, yet the resulting STAT5B-driven transcriptional responses vary based on instigating cytokine and/or differentiation state of responding cells.

**Fig. 4.**
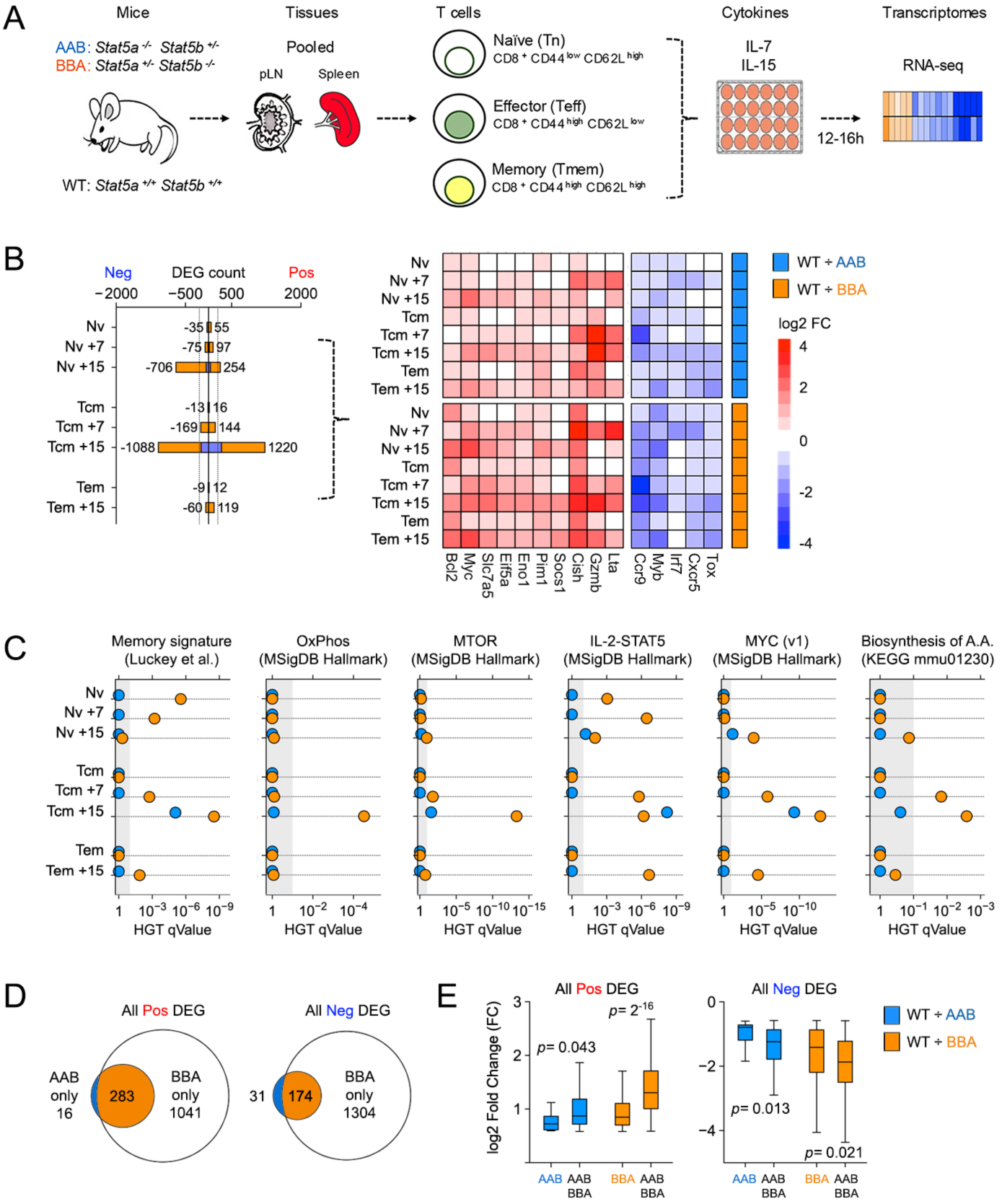
STAT5 paralog asymmetry in CD8^+^ T cells. (A) Cartoon outlines experimental system for transcriptome studies. (B) Stacked bar plot enumerates Differentially Expressed Genes (DEG) in KO T cells relative to WT controls (blue stack = AAB versus WT; orange stack = BBA versus WT). Heat map shows log2 fold change values for representative DEG across genotypes, cytokines and cell states. (C) Positively regulated DEG were subjected to hypergeometric testing against the indicated databases. Scatter plots show enrichment *q* values for top STAT5-regulated pathways across genotypes, cytokines and cell states (blue = AAB versus WT, orange = BBA versus WT). (D) Positively (left) and negatively (right) regulated DEG were compiled across cytokines and cell states. Venn plot compares DEG from AAB and BBA cells. (E) Box plots show log2 fold change values for DEG classes identified in (D). AAB = DEG called only in AAB cells, BBA = DEG called only in BBA, AAB/BBA = DEG called in both genotypes. Blue = pairwise comparison of AAB versus WT, orange = BBA versus WT. All DEG sets listed in Table S1 and usage detailed in Table S2. Ex vivo culture conditions detailed in Table S7. Replicate counts and statistical tests for all experiments are detailed in Table S6.

Next, we compared DEG called across cell states and stimuli to determine if any are solely dependent on STAT5A or STAT5B. This analysis revealed that almost every DEG called in AAB cells was also called in BBA cells, indicating that few (if any) are strictly STAT5A dependent (Fig. 4D). On the other hand, the vast majority of DEG called in BBA cells were not called in AAB cells, suggesting that these are STAT5B dependent (Fig. 4D). However, since STAT5B twice as abundant (Fig. 2E-F), it is also possible (if not likely) that these DEG reflect differences in total STAT5 content rather than *bona fide* preference for STAT5B. Building on this latter point, we also noted that the degree of induction (i.e. fold change values) differed between ‘overlapping’ and ‘unique’ DEG, the former appearing more labile (Fig. 4E). Thus, we interpret that ‘high mobility’ target genes employ both STAT5A and STAT5B, while ‘low mobility’ targets genes mainly employ STAT5B because STAT5A is less available. Immunologically relevant examples of low mobility STAT5 target genes include *Bax*, *Bhlhe40* and *Nfil3* (i.e. DEG with average fold change <1.3 across STAT5B datasets; Table S3).

### STAT5 paralog redundancy in CD8^+^ T cells

Upon activation, STAT5A and STAT5B distribute through the genome via DNA binding motifs known as “Gamma interferon Activated Sequences” (GAS elements)(3). To determine if they co-localize, we compared genome-wide distributions using ChIP-seq datasets generated in effector CD8^+^ T cells stimulated with IL-2 (7). Crucially, these datasets were chosen because absolute numbers of STAT5A- or STAT5B-bound regions, colloquially termed ‘peaks’, are similar (Fig. 5A). We began by identifying peaks enriched over background, then merged those within 100 base pairs of one another to minimize ‘satellites’ that can flank high amplitude peaks (Fig. 5A, Table S4). Next, we cross-referenced these merged peak sets to identify genomic regions bound by STAT5A and/or STAT5B. Based on prior studies (7, 33), we expected that most captured regions would be bound by both and, indeed, that is what we found. Approximately 2/3 of STAT5A peaks overlapped with STAT5B peaks (and vice versa), while the remaining 1/3 appeared unique to STAT5A or STAT5B (Fig. 5A). We then compared peak summit values and learned that regions bound by both STAT5A and STAT5B had greater means and contained all of the highest amplitude events, suggesting these are robust, high mobility target sites (Fig. 5B). DNA motif analysis also showed that peaks bound by both are far more likely to bear STAT binding motifs than those bound by one or the other (Fig. 5C). Thus, STAT5A and STAT5B tend to co-localize, especially at high traffic sites that are likely to have functional consequence.

**Fig. 5.**
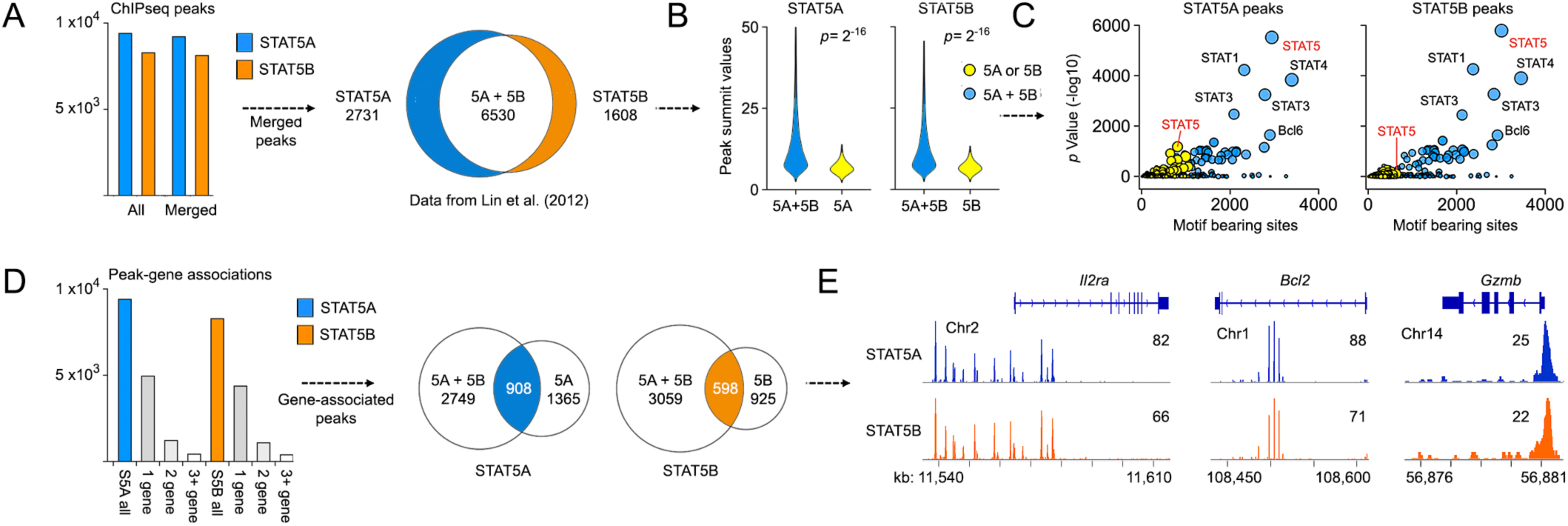
STAT5 paralog redudancy in CD8^+^ T cells. (A) Bar plot enumerates DNA regions bound by STAT5A (blue) or STAT5B (orange) pre- and post-peak merge. Venn plot compares merged regions. Union indicates at least 1 base pair overlap between STAT5A- and STAT5B-bound regions. (B) Violin plots compare peak summit values (i.e. peak amplitude) for STAT5A (left) and STAT5B (right) bound regions across categories defined in (A). 5A = regions bound only by STAT5A, 5B = regions bound only by STAT5B, 5A+5B = regions bound by STAT5A and STAT5B. (C) Scatter plot shows total motif bearing sites and associated *p* values for STAT5A (left) or STAT5B (right) bound regions. Color coding as in (B). Point size is proportional to *p* value. (D) STAT5A and STAT5B bound regions were assigned to genes based on linear proximity. Bar plot enumerates those assigned to one or more genes. Venn plot compares Peak-Associated Genes (PAG) across categories defined in (A). 5A = PAG associated with only STAT5A, 5B = PAG associated with only STAT5B, 5A+5B = PAG associated with STAT5A and STAT5B. (E) Genome browser histograms show mirroring of STAT5A and STAT5B bound regions at select loci. All STAT5A and STAT5B bound regions are catalogued in Table S4. Replicate counts and statistical tests for all experiments are detailed in Table S6.

To infer transcriptional effects, we next assigned peaks to genes based on genomic proximity. As expected, just over half could be unambiguously assigned, with the majority mapping to a single gene (Fig. 4D). Next, we compared Peak Associated Genes (PAG) and found that, like peaks, the vast majority of PAG were associated with both STAT5A and STAT5B (Fig. 4D). This was plainly evident at emblematic STAT5 target genes like *Il2ra*, *Bcl2* and *Cish*, where STAT5A and STAT5B peaks closely mirrored (Fig. 4E). We also noted a substantial number of PAG associated either STAT5A or STAT5B but, upon inspection, determined that most of these were not strictly dichotomous. Typically, when a single paralog was assigned to a gene, associated peaks were only slightly under the chosen *p* value threshold, while peaks for the other paralog were only slightly over (Fig. S4). Furthermore, PAG associated with either STAT5A or STAT5B tended to be lower amplitude and contained less STAT motif enrichment, again suggesting that they are not robust targets (Fig. 5B-C).

To further explore redundancy between STAT5 paralogs, we used restored STAT5A in STAT5A-deficient CD8^+^ T cells, then measured transcriptomes by RNA-seq, reasoning that if *bona fide* STAT5A-specific genes exist, then constitutively active STAT5A (CA-STAT5A) should selectively mobilize these and not putative STAT5B-specific genes (Fig. 6A). For these experiments, naïve CD8^+^ T cells were sorted from AAB mice, stimulated with agonist anti-TCR and anti-CD28 antibodies, then transduced with a retroviral vector expressing constitutively active STAT5A (CA-STAT5A). DEG were called relative to cells transduced with ‘empty’ control vector. Strikingly, hundreds of DEG were detected, including many well-known STAT5 targets like BCL2 and GZMB (Fig. 6A). Next, we cross-referenced with ChIP-seq data to segregate DEG based on whether they are proximally bound by STAT5A and STAT5B (5A + 5B), STAT5A alone (5A only), STAT5B alone (5B only) or neither (No STAT5)(as per Fig. 5D). Of note, the chosen ChIP-seq dataset was generated in CD8^+^ T cells cultured *in vitro* with IL-2 (7), similar to our retroviral system. Surprisingly, we of found that most DEG were not engaged by STAT5A or STAT5B, suggesting that much of the downstream transcriptional response in this setting is indirect (Fig. 6B). However, direct effects were also evident; 182 DEG were bound by both STAT5A and STAT5B, while 39 were bound solely by STAT5A (Fig. 6B). It is tempting to classify the latter as STAT5A-specific, but we hesitate to do so because a similar number of DEG were engaged solely by STAT5B (Fig. 6B). Also, DEG bound solely by STAT5A or STAT5B had lower fold-change values than those bound by neither or both (Fig. 6C-D), and no meaningful pathway enrichment was detected among STAT5A and STAT5B ‘only’ DEGs (not shown), unlike ‘shared’ DEG, which were highly enriched for known STAT5-regulated pathways like JAK-STAT signaling and apoptosis (Fig. 6E). Thus, it may be some loci are better or even exclusively engaged by STAT5A or STAT5B, but the bulk of evidence suggests that few (if any) *bona fide* STAT5 targets are solely and strictly dependent on one or the other.

**Fig. 6.**
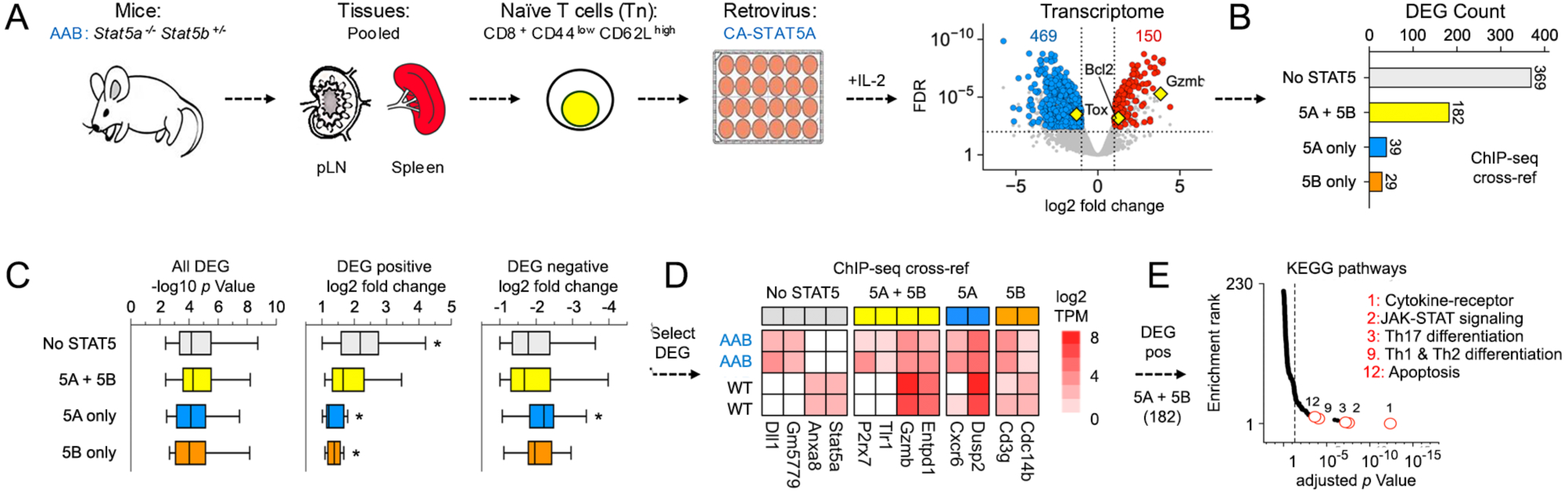
STAT5 paralog specificity is not evident in CD8^+^ T cells. (A) *Stat5a^−/−^ Stat5b^+/−^* CD8^+^ T cells were transduced with CA-STAT5A, then assayed by RNA-seq. Cartoon outlines experimental system. Volcano plot shows positively (red) and negatively (blue) regulated DEG from pairwise comparison of CA-STAT5 and ‘empty’ control (both in AAB CD8^+^ T cells). (B) DEG were cross-referenced with STATA or STAT5B Peak-Associated Genes (PAG from Fig. 5D). Bar plot enumerates DEG associated with STAT5A only (5A), STAT5B only (5B), both (5A + 5B), or neither (no STAT5). (C) Box plots compile *p* values and log2 fold change values for DEG segregated as in (B). (D) Heat map shows log2 fold change values for representative DEG from each category in WT or AAB cells (each vs. ‘empty’ control). (E) DEG associated with both STAT5A and STAT5B (yellow bar; 182 total) were subjected to hypergeometric testing against the KEGG database. RankLine plot shows enrichment *p* values and ranks for all captured pathways. Select pathways are noted (rank shown). All DEG sets are catalogued in Table S1 and usage detailed in Table S2. Replicate counts and statistical tests for all experiments are detailed in Table S6.

### Evidence for cytokine, lineage and state restricted STAT5 activities

Having established that STAT5B is dominant in CD8^+^ T cells, we next compared STAT5B-driven transcriptomes across cytokines and differentiation states. First, we noted a striking distinction between cytokines; IL-15 mobilized nearly 10 times as many DEG as IL-7, regardless of differentiation state (Fig. 7A). We also found that the character of downstream responses differed; only 35% of DEG mobilized by IL-7 were also mobilized by IL-15 in naive cells (61 of 172), versus 81% (253 of 311) in memory cells (Fig. 7A-C, Table S5). Moreover, only 17% of DEG mobilized by IL-7 in naive T cells were also mobilized by IL-7 in memory cells (29 of 172), versus 65% for IL-15 (627 of 960)(Fig. S5A-C, Table S5). Thus, we find evidence for: (1) cytokine specificity (e.g. differences between IL-7 and IL-15 in Tn cells), (2) cell state specificity (e.g. differences between IL-7 in Tn and Tcm), and (3) stereotypical responses (e.g. similarities between IL-15 and IL-7 in Tcm, and IL-15 in Tn and Tcm). As such, we can infer that STAT5-driven transcriptional responses vary downstream of IL-7 and IL-15, and that the amount of variance depends on cellular state, with naïve cells exhibiting more pronounced differences than memory cells.

**Fig. 7.**
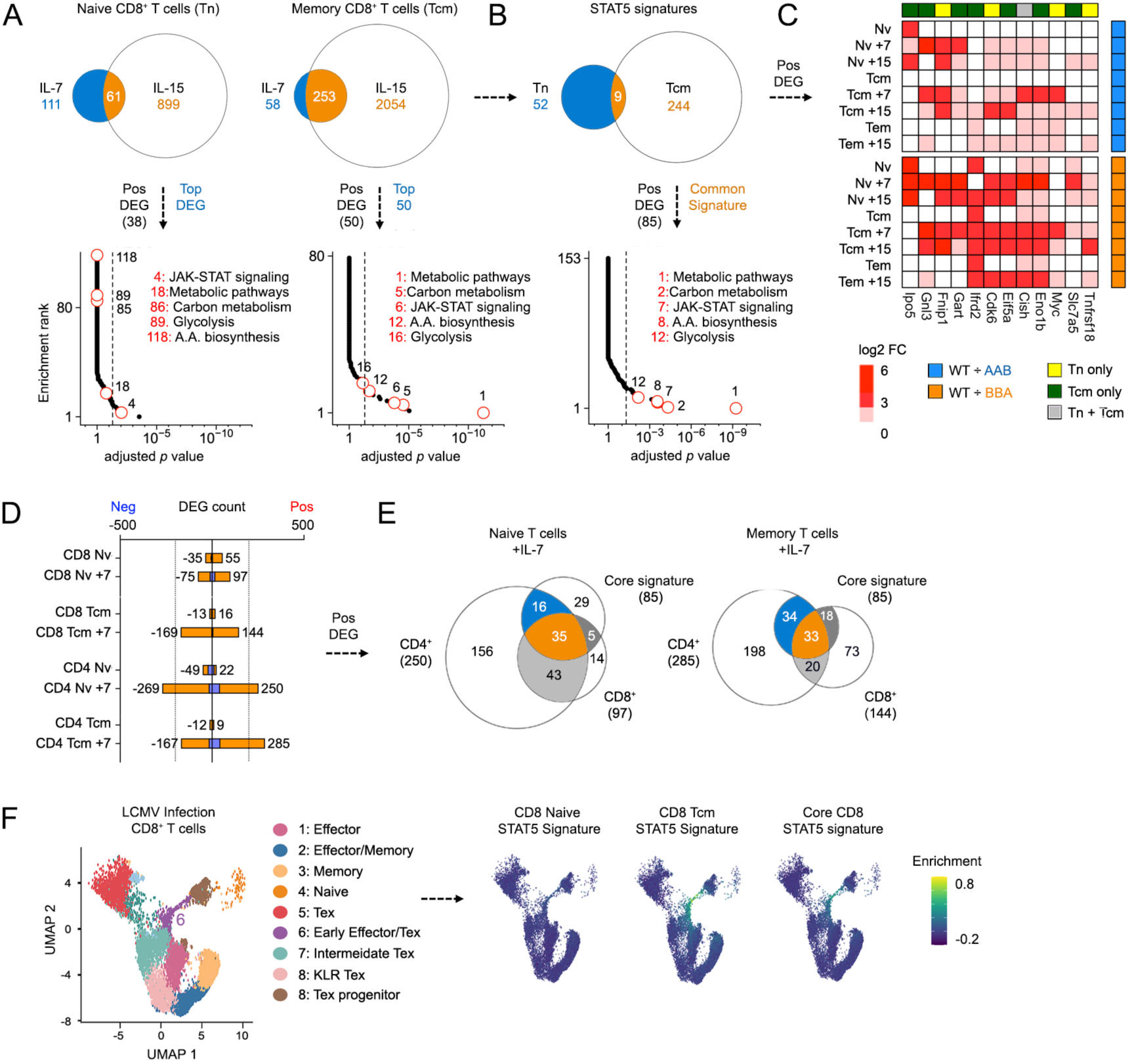
A core STAT5 signature for CD8^+^ T cells. (A) Venn plots compare DEG mobilized in naïve (left) or memory (right) CD8^+^ T cells (DEG from Fig. 4). Union represents genes mobilized by both IL-7 and IL-15 in each cell type. Positively regulated ‘union’ DEG from naïve (38 total) and memory (top 50) CD8^+^ T cells were also subjected to hypergeometric testing against the KEGG database. RankLine plots show enrichment *p* values and ranks for all captured pathways. Select pathways are noted (rank shown). (B) Venn plot compares ‘union’ DEG from (A). RankLine plot shows HGT results for core STAT5 signature genes against the KEGG database. Select pathways are noted (enrichment rank shown). Core STAT5 signature combines all 38 positively regulated genes contained within the IL-7 union set and the top 50 positively regulated genes DEG from the IL-15 union set (ranked based on fold change). 3 genes are shared, so this yields an 85-element signature. (C) Heat map shows select genes contained in the core STAT5 gene signature (full roster shown in Fig. S5D). (D) Stacked bar plot enumerate Differentially Expressed Genes (DEG) in CD4^+^ and CD8^+^ T cells relative to WT controls (blue stack = AAB versus WT; orange stack = BBA versus WT; CD8^+^ T cell data same as Fig. 4). (E) Left Venn plot compares core STAT5 signature to DEG positively regulated by IL-7 in naïve CD4^+^ or CD8^+^ T cells. Right Venn plot compares to memory CD4^+^ or CD8^+^ T cells. (F) scRNA-seq UMAP projection segregates CD8^+^ T cells responding to acute or chronic LCMV infection. Clustering and annotation are as published. Feature plots show module score enrichment for the indicated gene signatures. All gene signatures catalogued in Table S5. Replicate counts and statistical tests for all experiments are detailed in Table S6.

To determine if lineage identity also impacts STAT5B activity, we compared transcriptomes from CD4^+^ and CD8^+^ T cells (Fig. 7D-E, Table S1-S2). These studies focused exclusively on IL-7 because IL-7R⍺ is similarly expressed on CD4^+^ and CD8^+^ T cells, unlike IL-15 receptors components IL-2Rβ and IL-15R⍺, which are more abundant CD8^+^ T cells. First, we performed RNA-seq on naive and memory CD4^+^ T cells sorted from LNs and spleens of WT, AAB or BBA mice (Fig. 7D, Fig. S5E). Then, we called DEG and cross-referenced with corresponding CD8^+^ T cell datasets to learn that: (1) IL-7 mobilizes more DEG in CD4^+^ T cells than in CD8^+^ T cells (Fig. 7D), (2) most DEG and pathways mobilized by IL-7 in naive CD8^+^ T cells are also mobilized in naïve CD4^+^ T cells (132 of 172 = 76%; Fig. S5F-H), and (3) relatively few DEGs and pathways mobilized by IL-7 in memory CD8^+^ T cells were also mobilized in memory CD4^+^ T cells (92 of 313 = 29%; Fig. S5G-H). Thus, STAT5 activity does vary across T cell lineages, and, again, the amount of variance depends on cellular state although, here, memory cells exhibit more than naïve cells.

### A core STAT5 signature for CD8^+^ T cells

Next, we sought to define a core STAT5 signature evident across cytokines and differentiation states. To begin, we compared all DEG called in naive and memory CD8^+^ T cells and found that they are strikingly distinct; only 15% overlapped (9 of 61; Fig. 7B, Table S5). That dichotomy was also evident at pathway level as naive cells were highly enriched for JAK-STAT signaling while memory cells were highly enriched for metabolic pathways (Fig. 7A). Next, we devised a composite, or ‘core’, signature by merging pan-cytokine DEG from naive and memory cells, focusing on those that are positively regulated (i.e. those induced by STAT5-activating cytokines; Fig. 7B-C; Fig. S5D, Table S5). Predictably, pathway analysis reflected the composite nature of this gene set; both JAK-STAT signaling and metabolic pathways were highly enriched (Fig. 7B).

Several genes in our core signature are known to be induced by STAT5 and upstream cytokines in CD4^+^ T cells. Notable examples include *Il2ra*, *Myc* and *Slc7a5* (Fig. 7C, Fig. S5D)(6). To determine if our core STAT5 signature can distinguish between CD4^+^ and CD8^+^ T cells, we cross-referenced the pan-cytokine CD8^+^ T cell DEG set from Fig. 7A with naïve or memory CD4^+^ T cell DEG sets from Fig. 7D. The results was clear: a majority of core signature elements are contained within the CD4^+^ T cell gene sets (51/85 = 60% for naïve cells, 67/85 = 79% for memory cells; Fig. 7E). Therefore, despite the fact that it was built exclusively from CD8^+^ T cell data, our core STAT5 signature likely cannot distinguish between STAT5 responses in naive CD4^+^ and CD8^+^ T cells.

To see if our core STAT5 gene signature can be used to probe for STAT5 activity, we mined a single-cell RNA-seq (scRNA-seq) dataset composed of CD8^+^ T cells from mice challenged with acute and chronic Lymphocytic Choriomeningitis Virus (LCMV) (57). After recreating the published UMAP (Fig. 7F), we performed ‘module score’ analysis to determine which areas (if any) are enriched for our core STAT5 signature and/or its constituents (Table S5). Strikingly, we found that signatures derived from memory CD8^+^ T cells were highly enriched, whether based on IL-7 alone, IL-15 alone or both cytokines (Fig. 7F and Fig. S6A-B). By contrast, those derived from naive CD8^+^ T cells were not enriched at cluster level (Fig. 7F and Fig. S6A), and individual events reflecting transient encounters with STAT5-activating cytokines were not detected (Fig. S6B). Also notable was the position of enrichment. When present, it localized to a UMAP region containing ‘early effector or exhausted’ cells which recently encountered antigen and, thus, are likely experiencing acute STAT5 activity downstream autocrine or paracrine IL-2 responses (cluster 6; Fig. 7F and Fig. S6A-B). Thus, our core signature and its constituents may be useful for detecting strong, synchronized STAT5 signaling associated with effector responses, but not weaker, asynchronous signaling associated with homeostatic responses.

## Discussion

The transcription factor STAT5 is fundamental to lymphocyte biology. However, the relationship between its two paralogs, STAT5A and STAT5B, and the extent to which they are functionally distinct, remains unclear. Here, we engage this longstanding question using a combination of genetic and genomic tools focused on CD8^+^ T cells, key players in pathogen-, tumor-, and self-driven inflammation. Previously, we used a similar approach to demonstrate that STAT5B is dominant over STAT5A in CD4^+^ T cells and ILCs, where it influences numerous developmental and effector pathways (30, 35). Here, we establish that the ‘STAT5B > STAT5A’ rule holds true across the lymphoid compartment and present evidence for a shared mechanism: STAT5B is twice as abundant in CD8^+^ T cells, like it is in CD4^+^ T cells and ILCs. These findings affirm the centrality of STAT5 in CD8^+^ T cells, advance basic understanding of why it is so important, and resolve the question of whether STAT5A and STAT5B are redundant or functionally distinct, concluding that asymmetric expression enables both classifications to co-exist.

Although overarching trends are similar (STAT5B> STAT5A), CD8^+^ T cells are clearly more sensitive to STAT5B-deficiency than CD4^+^ T cells. In fact, only CD8^+^ T cells are depleted in STAT5B-deficient mice, resulting in sharply skewed CD4/CD8 ratios. This finding is line with prior work on CD8^+^ T cells (33) and ILCs, which are closely related to T cells and also depleted in STAT5B-deficient mice (35,58). Here, it is important to emphasize that, unlike phenotypic differences between STAT5A- and STAT5B-deficient cells, asymmetric expression of STAT5A and STAT5B likely does not explain why CD8^+^ T cells are more sensitive to STAT5 deficiency than CD4^+^ T cells, given total STAT5 levels are similar across these lineages.

STAT5B-deficiency dramatically illustrates the importance of STAT5 signaling in CD8^+^ T cells. In fact, aberrant CD8^+^ T cell differentiation is a cardinal feature of STAT5B-deficient mice, which are characterized by loss of naive quiescence and a corresponding shift towards of effector- and memory states. Given the lack of acute stimulus (e.g. infection, immunization), we interpret that this reflects self-reactivity unleashed by defective Treg-mediated suppression, in line with the idea that Treg are impaired in STAT5B-deficient mice (30, 33). However, it is also worth noting that, while prone to effector/memory differentiation, STAT5-deficient CD8^+^ T cells cannot elaborate key elements of the cytotoxic effector program (e.g. granzyme expression). Thus, excessive killing likely does not contribute to the attendant kidney disease seen in STAT5B-deficient mice (30). Nevertheless, it is clear that STAT5B is critical for cytokine-driven gene expression in CD8^+^ T cells and that, in turn, many important cellular pathways are impacted by STAT5B-deficiency at transcriptional level, including MTOR, MYC and several involved with metabolism, befitting the key role of STAT5 in that arena (1, 6).

Consistent with prior work (33), we demonstrate that STAT5B-deficiency has far greater impact on gene expression in CD8^+^ T cells than STAT5A-deficiency. However, genome-wide analysis did not reveal major differences in the distribution of STAT5A and STAT5B, in line with the original report of this dataset (7) and recent ChIP-seq studies in CD8^+^ T cells (33). Instead, we and others have found that STAT5A and STAT5B mostly co-localize and, thus, have similar target ranges. This ‘mirroring’ is evident across the genome and most conspicuous at high amplitude sites decorating *bona fide* STAT5 targets, like *Il2ra* and *Gzmb*. Furthermore, using a gain-of-function system, we established that transcripts mobilized by STAT5A mainly come from loci engaged by both STAT5A and STAT5B, rather STAT5A alone. Thus, broadly speaking, STAT5A and STAT5B appear to engage the same target genes, making them redundant at molecular level. Still, we cannot fully discount the possibility that some genes are regulated solely by STAT5A or STAT5B. Indeed, we identified: (1) 47 transcripts whose expression was disturbed only in STAT5A-deficient cells, (2) 4339 genomic regions engaged only by STAT5A (25% of all peaks), and (3) 39 genes mobilized by STAT5A and engaged only by STAT5A. However, we also noted that: (1) DEGs called only in STAT5A- or STAT5B-deficient cells are ‘weaker’ than those called only in both (i.e. had lower fold change values), (2) ChIP-seq peaks for STAT5A- or STAT5B-specific regions are ‘weaker’ than those for shared regions (i.e. lower peak amplitudes, higher p values), and (3) a similar number of genes mobilized by STAT5A were engaged only by STAT5B. Therefore, the bulk of evidence suggests that few (if any) genes are controlled solely by STAT5A or STAT5B and that if such genes exist, transcriptional output is likely lesser than that of genes subject to both.

Given STAT5B emerged as the dominant paralog, we compared STAT5B-driven transcriptomes across cytokines, lineages and differentiation states. This analysis revealed that downstream consequences vary based on the instigating cytokine (IL-7 or IL-15), and that the scale of this variance depends on differentiation state, with naïve cells exhibiting more pronounced differences than memory cells. A similar trend emerged when comparing CD4^+^ and CD8^+^ T cells; downstream consequences vary based on lineage identity and, again, the scale of variance depends on cellular differentiation state. Thus, we present evidence for both cytokine specific STAT5 activities (e.g. differences between IL-7 and IL-15 in Tn cells) and lineage specific STAT5 activities (e.g. differences between IL-7 in CD4^+^ and CD8^+^ T cells).

We also devised a core signature comprised of STAT5B regulated genes evident across cytokines and differentiation states. Importantly, this core signature includes only positively regulated genes (i.e. genes induced by IL-7 or IL-15 in a STAT5-dependent manner) because unimodal genesets perform better than multimodal genesets when subjected to Hypergeometric Testing (HGT), the standard methodology for geneset/pathway analysis. We also selected the ‘top 50’ from the IL-15 geneset to approximate the 38 positively regulated genes contained within the IL-7 geneset, and because HGT performs best with genesets ranging from 50-200 elements. To assess the utility, we probed a single-cell RNA-seq dataset which captures CD8^+^ T cells responding to LCMV infection, a gold standard model for cytotoxic T cell responses (59). Remarkably, we found that our core signature was enriched in only one cluster of cells annotated as ‘early effectors’. We also learned that this enrichment was driven mainly by signature elements derived from memory and IL-15 genesets rather than naive and IL-7 genesets. Neither the core STAT5 signature nor any of its constituent genesets were enriched in the ‘naive’ or ‘exhausted’ clusters, apart from ‘intermediate exhausted’ cells. This is an intriguing result as it suggests that STAT5 signaling is progressively disabled as cells move from early (or intermediate) to terminal exhaustion. Notwithstanding, it is evident that our core signature and its constituents can be useful for detecting strong synchronized STAT5 responses associated with early T cell activation, as in early effectors, but not weaker homeostatic STAT5 responses, as in naive or resting memory T cells. These data again affirm the central role of STAT5 in CD8^+^ T cells and spotlight acute, synchronized cytokine signaling as a driving force for effector and memory CD8^+^ T cell responses.

## Methods

### Experimental animals

STAT5 ‘allele’ mice were generated as described (30, 35, 39). Briefly, mice lacking the entire STAT5 locus on one chromosome (*Stat5a/b^+/−^*) were crossed with mice lacking one allele of STAT5A (*Stat5a^+/−^ Stat5b^+/+^*) or STAT5B (*Stat5a^+/+^ Stat5b^+/−^*) to produce 8 allele combinations (Fig. 1A). We refer to each strain according to the alleles that were deleted, except for wild type controls (*Stat5a^+/+^ Stat5b^+/+^*). Thus, AAB mice lack two alleles of STAT5A and one of STAT5B (*Stat5a^−/−^ Stat5b^+/−^*; always colored blue), while BBA mice lack two alleles of STAT5B and one of STAT5A (*Stat5a^+/−^ Stat5b^−/−^*; always colored orange). *Stat5a/b* ^flox/flox^ *Cd4-* Cre*^+/−^* mice were generated as described (5). *γc^−/−^ Rag2^−/−^* mice were purchased from Taconic Farms. Both male and female mice were used and sex matched within experiments. Animals were housed and handled in accordance with NIH guidelines and all experiments approved by either the NIAMS or University of Miami Animal Care and Use Committee.

### Cell purification and culture

Tissues were dissected from 8-16 week old mice and processed to single cell suspensions. Mesenteric lymph nodes (mLN) and peripheral lymph nodes (pLN; inguinal, brachial, axillary and superficial cervical lymph nodes) were mechanically dissociated through 40 uM cell strainers. Spleens were first mechanically dissociated, then red blood cells depleted by hypotonic lysis (Gibco/Life Technologies). Bone marrow was flushed from rear femurs and red blood cells depleted by hypotonic lysis. CD4^+^ or CD8^+^ T cells were enriched by magnetic separation (Miltenyi kits) or cell sorting (as below). *Ex vivo* culture conditions for all experiments are detailed in Table S7.

### Cytometry

For surface antigens, cells were stained and washed in phosphate-buffered saline supplemented with 0.5% bovine serum albumin and 0.1% sodium azide directly *ex vivo*. For intracellular antigens (Fig. 3D), naive cells (CD3^+^, CD8⍺^+^, CD44 ^low^, CD62L ^high^) were sorted from pooled LN and spleens, stimulated with plate-bound anti-CD3ε and anti-CD28 (1 μg/ml each) in the presence of human IL-2 (100 U/ml; NIH/NCI BRB Preclinical Repository) for 48 hours, then fixed and permeabilized with Cytofix/Cytoperm (BD Biosciences). Fluorochrome labelled antibodies were purchased from Thermo-Fisher, BD Biosciences or Biolegend. Dead cells were excluded using Live/Dead Aqua (Invitrogen) or Zombie NIR (Biolegend). Total STAT5 protein was measured in splenocytes directly *ex vivo*. Cells were fixed with 2% formaldehyde, permeabilized with 100% methanol, then stained with a rabbit polyclonal IgG that recognizes both STAT5A and STAT5B (sc-835; Santa Cruz Biotechnology, Santa Cruz, CA) in conjunction with Phycoerythrin labelled goat anti-rabbit IgG for detection (ac-3739; Santa Cruz Biotechnology). Normal rabbit IgG was used as a negative control (ac-2027; Santa Cruz Biotechnology).

### Bone Marrow Chimeras

Lineage positive cells were depleted from WT (CD45.1 or CD90.1 congenic), *Stat5a^−/−^ Stat5b^+/−^* (AAB) or *Stat5a^+/−^ Stat5b^−/−^* (BBA) bone marrow by negative selection using ‘Lineage Cell Depletion Kit’ supplemented with biotinylated anti-NKp46 and anti-CD25 (Miltenyi Biotec). Cells were then washed and re-suspended in PBS before WT and KO cells were mixed (1:5 WT to KO ratio) and intravenously injected to sex-matched *γ*c−/− *Rag2*−/− hosts (2 x 10^5^ total cells per mouse in 300 ul PBS). Engraftment was measured 8-12 weeks later in spleen.

### RNA-seq

Naive (CD3^+^, CD4^+^ or CD8⍺^+^, CD44 ^low^, CD62L ^high^), central memory (CD3^+^, CD4^+^ or CD8⍺^+^, CD44 ^high^, CD62L ^high^) and effector memory (CD3^+^, CD8⍺^+^, CD44 ^high^, CD62L ^low^) T cells were sorted (>95% purity) from pooled LN and spleens, then either processed directly *ex vivo* or cultured for 12-18 hours with IL-7 or IL-15 (10 ng/ml each; R&D Systems). *Ex vivo* culture conditions are further detailed in Table S7. 2-4 biological replicates were collected for each experimental group with similar numbers of cells per replicate (20-100 x 10^3^, depending on group/condition). These were lysed in Trizol reagent and total RNA purified by phenol-chloroform extraction with GlycoBlue as co-precipitant (7-15 μg per sample; Life Technologies). Poly(A)+ mRNA was then enriched by oligo-dT-based magnetic separation and single-end read libraries prepared with NEBNext Ultra RNA Library Prep Kit (New England Biolabs). Sequencing was performed with HiSeq 2500 (Illumina), then 50 bp reads (20-50 x 10^6^ per sample) aligned to mouse genome build mm10 with *tophat2*, assembled with *cufflinks* and gene-level counts compiled with *htseq-count*. To minimize normalization artifacts, genes failing to reach an empirically defined count threshold were purged using *htsfilter*. 12-14 x 10^3^ genes were typically recovered post filtering, regardless of genotype or experimental group. Counts were normalized and differentially expressed genes (DEG) called by quasi-likelihood F testing using *edgeR*. DEG call denotes >1.5 fold pairwise change, and Benjamini-Hochberg (BH) adjusted *p value* < 0.05. Transcripts per million (TPM) were compiled with *edgeR*. *clusterprofiler* was used for hypergeometric testing (HGT) against the KEGG, GO, Reactome, Molecular Signatures (MSigDB) databases, or custom genesets (catalogued in Table S1 and Table S5; usage detailed in Table S2). Heatmaps were rendered with pheatmap and all other plots with *ggplot2* or *Datagraph* (Visual Data Tools Inc.).

### ChIP-seq

STAT5A and STAT5B ChIP-seq datasets were downloaded from the NCBI sequence read archive via GSE36882 (7). Reads were aligned using bowtie, then non-redundant reads mapped to mouse genome mm9 using *macs2* with ‘input’ controls as reference for peak calling. *homer* was used to annotate peaks and test for TF motif enrichments. Gene proximal peaks were defined as occurring within introns, exons, UTRs or <10 kb of transcriptional start sites. Genome browser files were rendered with *IGV*. One of two biological replicates was used for all analyses.

### Retroviral transduction

Mutant mouse *Stat5a* (CA-STAT5A = S711F + H299R) was cloned into an MoMLV-based plasmid vector (MigR1) immediately before a dual internal ribosome entry sequence (IRES) and GFP cassette (6). CA-STAT5A plasmid and pCL-Eco ‘helper’ plasmid were then co-transfected into 293T cells (ATCC) using Lipofectamine (Invitrogen) and virus-containing supernatants collected 48 hours later. For transductions, CD8^+^ T cells were stimulated with plate-bound anti-CD3ε and anti-CD28 (10 μg/ml each) for 24 hours, exposed to viral supernatant for 1 hour (at 2200 rpm, 18°C), then cultured for a further 48 hours in the presence of human IL-2 (100 U/ml). Viable GFP^+^ cells were then sorted and processed for RNA-seq.

### scRNAseq

scRNA-seq gene expression (GEX) count matrixes were downloaded from NCBI via GSE188670 (59). Downstream analysis was performed with *seurat* using default function parameters, unless otherwise noted. Briefly, ribosomal transcripts were purged, then datapoints excluded if they had:>10% mitochondrial transcripts, >15,000 total transcripts, > 4,000 genes per cell, or <500 genes per cells. Next, individual samples were merged to a single dataset which was normalized and scaled using *scTransform*. Uniform Manifold Approximation and Projection (UMAP) reduction was then re-created using published XY coordinates at an empirically defined clustering resolution of 0.2. Cluster-defining markers were identified using *FindAllMarkers*. Feature and violin plots were rendered with *seurat*. *clusterprofiler* was used for pathway analysis, specifically hypergeometric testing of the top 400 markers enriched within each cluster (i.e. genes enriched in a given cluster relative to all others) against custom STAT5 signature geneSets (Table S5),

### Statistics

Statistical variances and distributions were measured by paired *t* test or Kolmogorov–Smirnov test per Table S6. Bonferroni correction was used to account for multiple testing in RNA-seq, ChIP-seq and pathway analysis datasets. Biological replicates for each experiment are detailed in Table S6. When present, error bars denote standard deviation across >2 biological replicates.

## Supporting information

Ristin_et_al_Sup_Tables

## Acknowledgments

We thank members of the O’Shea, Villarino and Malek labs for discussions, Gustavo Gutierrez-Cruz for sequencing support, and the NIAMS Flow Cytometry Group for cell sorting. Research in this publication was performed with support from the Flow Cytometry Shared Resource (SCR_022501) and Onco-Genomics Shared Resource (SCR_022502) of the Sylvester Comprehensive Cancer Center at the University of Miami Miller School of Medicine, which is supported by a National Cancer Institute Cancer Center Support Grant (CCSG) P30-CA240139 and State of Florida Bankhead Coley Research Infrastructure Grant 8BC09.

## Author contributions

Conceptualization: AVV Methodology: AVV Investigation: AVV, SR, AMF Data analysis: MD, CA, NI, LN Visualization: AVV, MD, LN Funding acquisition AVV, JOS Project administration: AVV Supervision: AVV, JOS, LH Writing – original draft: AVV Writing – review & editing: SR, MD, LH, AVV, JOS

## Data and materials availability

All data needed to evaluate the conclusions in the paper are present herein and/or the Supplementary Materials. For RNA-seq, gene-level transcript count tables are deposited to GEO under accession number. All analysis pipelines and code are available upon request.

## Supplementary Materials

6 supplementary figures, 7 supplementary tables:

**Supplementary table 1.** Gene set catalogue

**Supplementary table 2.** Gene set usage

**Supplementary table 3.** Low amplitude DEG

**Supplementary table 4.** ChIP-seq peak catalogue

**Supplementary table 5.** STAT5 signature

**Supplementary table 6.** Statistics catalogue

**Supplementary table 7.** Ex vivo culture conditions

## Supplemental Figure Legends

**Fig. S1.**
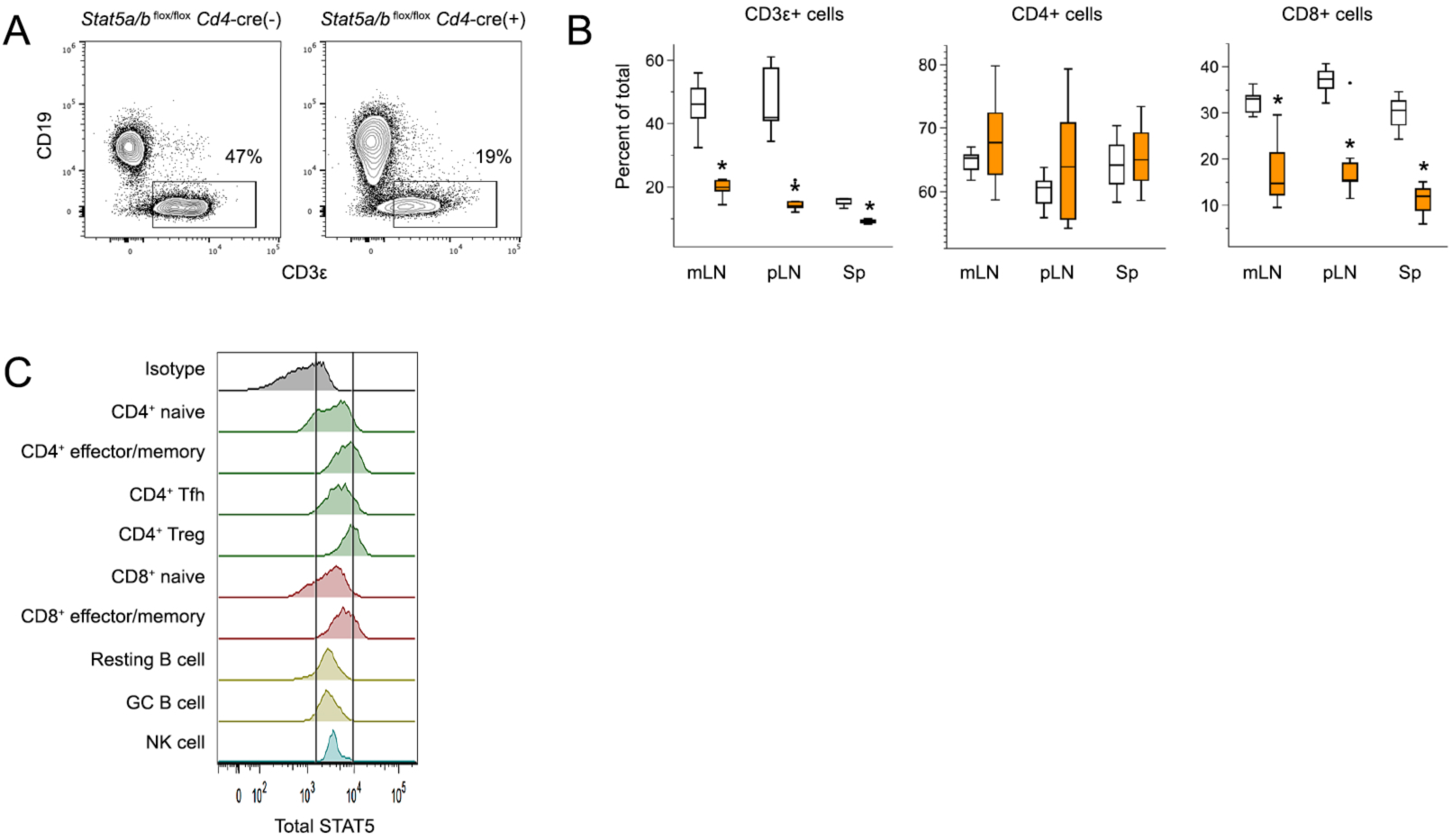
STAT5 deficiency markedly impacts CD8^+^ T cells. (A) Flow cytometry contour plots show surface CD3ε and CD19 proteins on live cells from pLN of *Stat5a/b ^flox/flox^ Cd4*-Cre^+/−^ mice and *Cd4*-Cre^−/−^ littermate controls. (B) Box plots compile frequencies of CD3ε^+^ T cells, CD3ε^+^ CD4^+^ T cells and CD3ε^+^ CD8^+^ T cells in mLN, pLN and spleens. (C) Flow cytometry histograms show total STAT5 protein levels across the lymphoid compartment in spleens of WT mice. Replicate counts and statistical tests for all experiments are listed in Table S6.

**Fig. S2.**
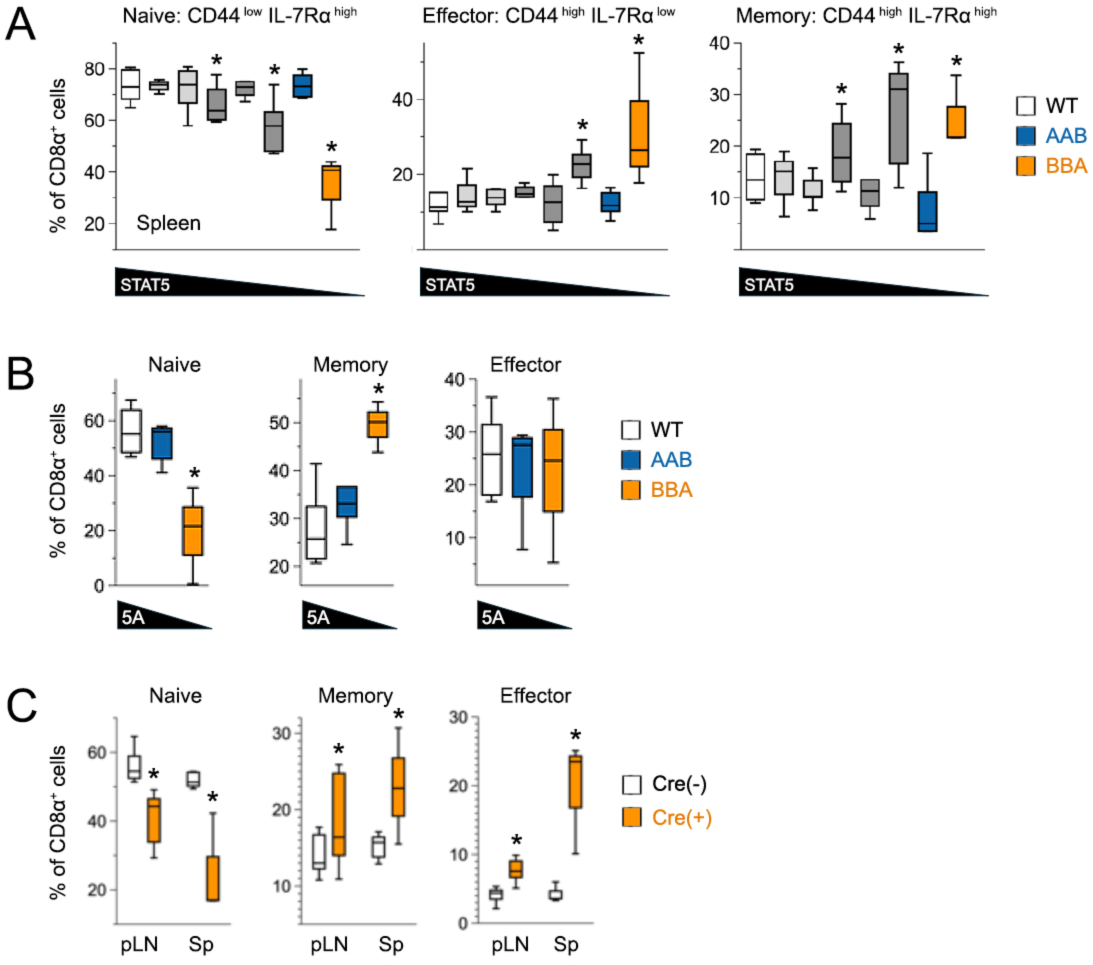
Outgrowth of effector and memory CD8^+^ T cells in STAT5-deficient mice. (A-B) Box plots compile frequencies of naïve, memory and effector CD8^+^ T cells in (A) spleens or (B) bone marrow of STAT5 ‘allele’ mice. (C) Box plots compile frequencies of naïve, memory and effector CD8^+^ T cells in pLN and spleens of *Stat5a/b ^flox/flox^ Cd4*-Cre^+/−^ mice and *Cd4*-Cre^−/−^ littermate controls. Replicate counts and statistical tests for all experiments are listed in Table S6.

**Fig. S3.**
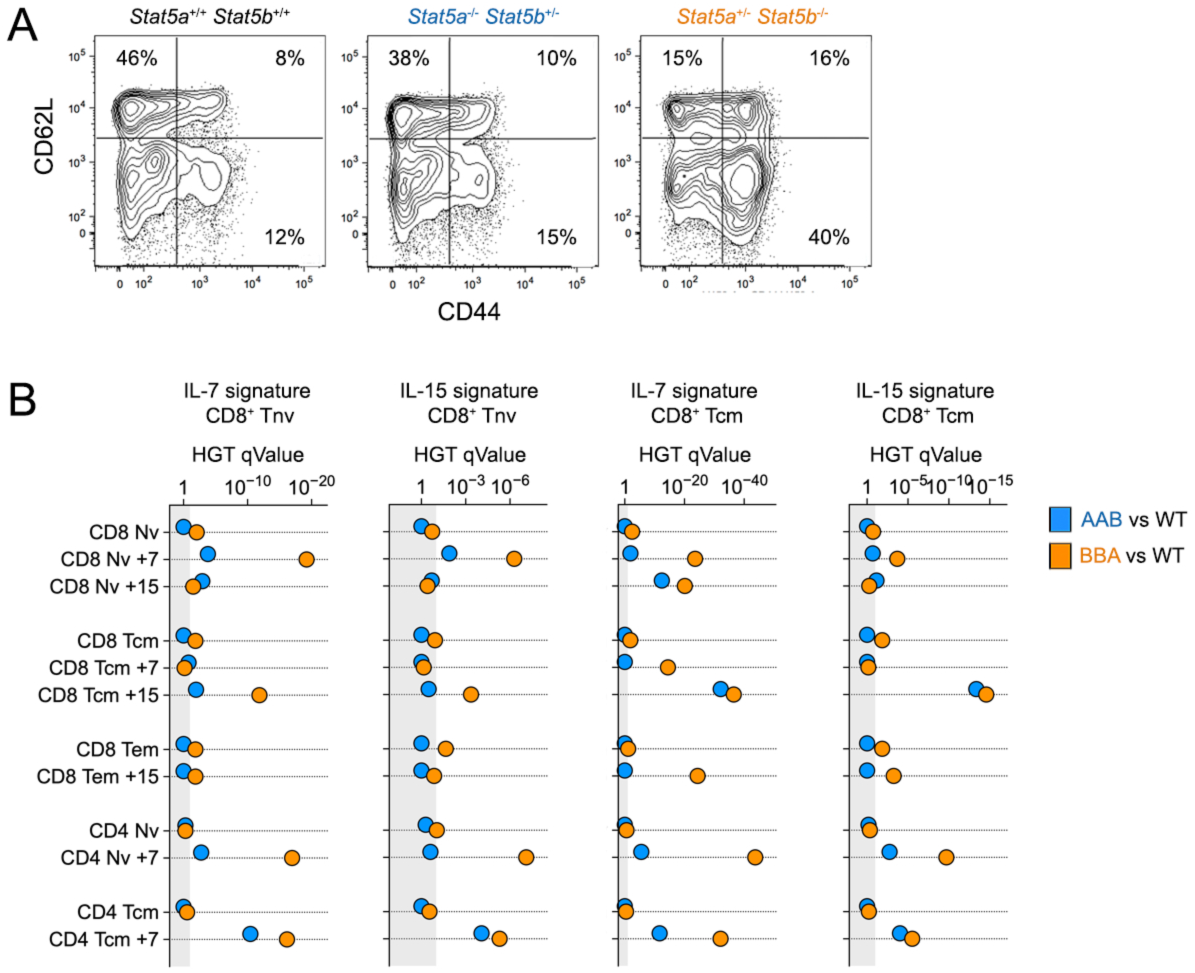
Disparities between STAT5A and STAT5B deficient CD8^+^ T cells. (A) Flow cytometry contour plots show surface CD44 and CD62L on CD8^+^ T cells from pLN. Naïve cells are defined as CD44^low^ CD62L^high^ (upper left), central memory cells as CD44^high^ CD62L^high^ (upper right) and effector cells as CD44^high^ CD62L^low^ (lower right). These were the markers and definitions used to sort CD4^+^ and CD8^+^ cells for RNA-seq (Fig. 4 and Fig. 7). (B) Positively regulated DEG were subjected to hypergeometric testing against the indicated databases (per Fig. 4C). Scatter plot shows enrichment q values for cytokine regulated genes across genotypes, cytokines and cell states (Y axis; blue = AAB versus WT, orange = BBA versus WT). All gene sets are catalogued in Table S1 and usage detailed in Table S2. Ex vivo culture conditions detailed in Table S7. Replicate counts and statistical tests for all experiments are listed in Table S6.

**Fig. S4.**
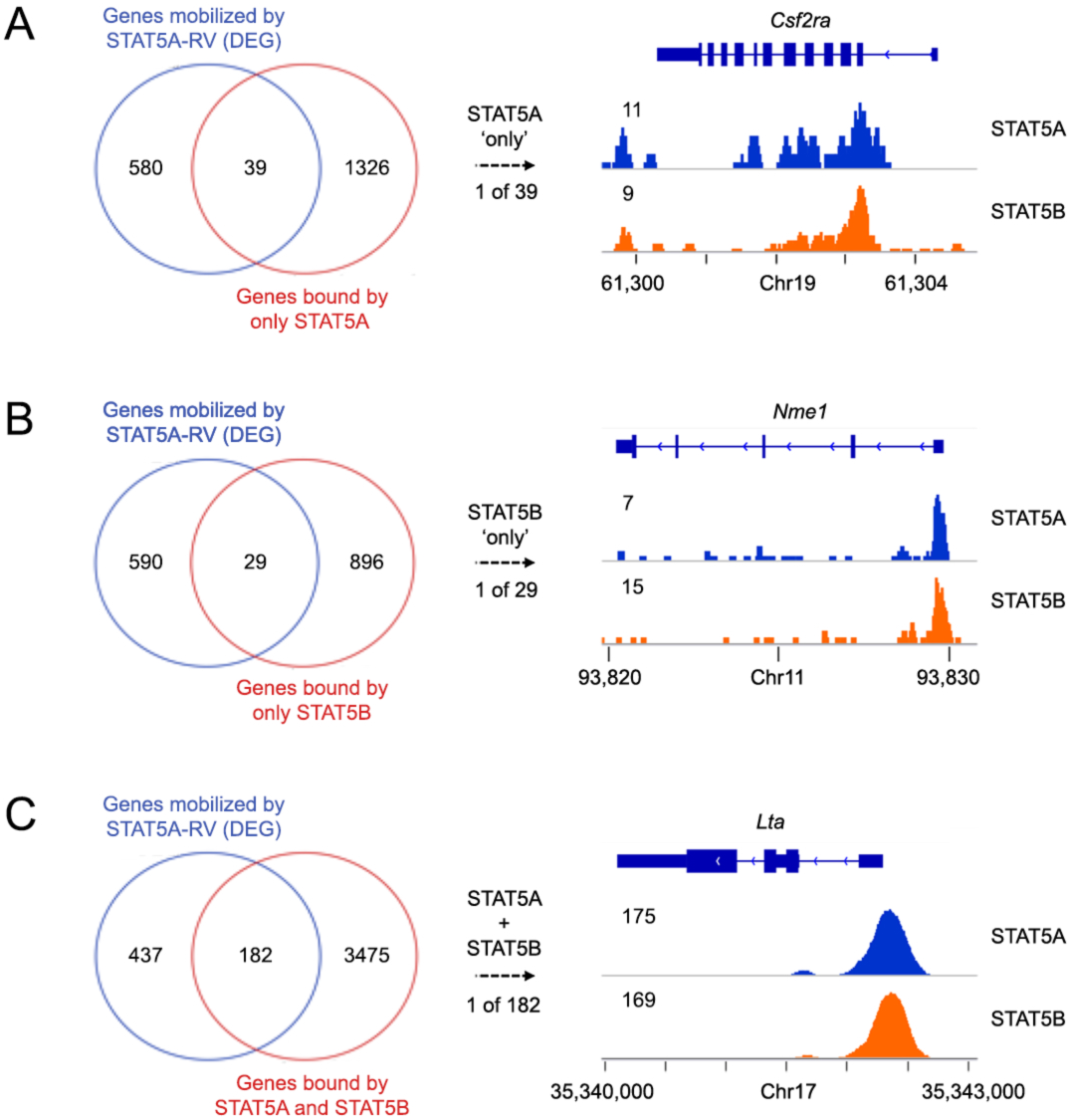
Genes bound only by STAT5A or STAT5B are not paralog specific. (A-C) Venn plots compare DEG mobilized by CA-STAT5A in STAT5A-deficient CD8^+^ T cells (DEG from Fig. 6) to Peak-Associated Genes (PAG) linked to (A) STAT5A alone, (B) STAT5B alone or (C) both STAT5A and STAT5B (PAG from Fig. 5D). Genome browser histograms show representative examples. All STAT5A and STAT5B bound regions are catalogued in Table S4. All gene sets are catalogued in Table S1 and usage detailed in Table S2. *Ex vivo* culture conditions for retroviral transduction are detailed in Table S7. Replicate counts and statistical tests for all experiments are listed in Table S6.

**Fig. S5.**
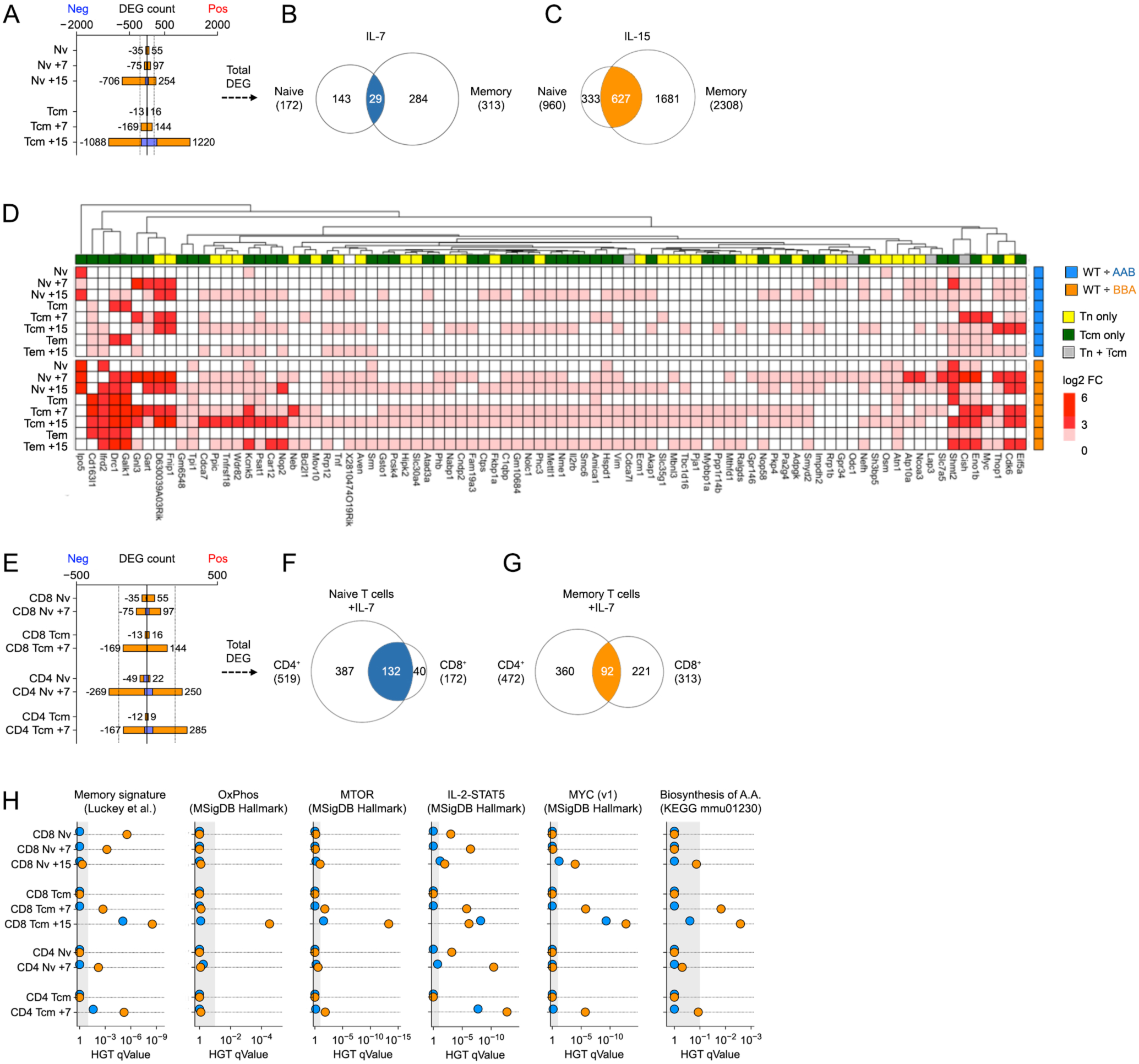
Core STAT5 signature assembly and validation. (A) Stacked bar plot enumerates DEG in KO CD8^+^ T cells relative to WT controls (blue stack = AAB versus WT; orange stack = BBA versus WT; DEG same as Fig. 4). (B-C) Venn plots compare DEG mobilized by (B) IL-7 or (C) IL-15 in naïve and memory CD8^+^ T cells. (D) Heat map shows log2 fold change values for all elements of the core STAT5 signature across genotypes, cytokines and cell states. (E) Stacked bar plot enumerates DEG in KO CD4^+^ or CD8^+^ T cells relative to WT controls (same as Fig. 7D). (F-G) Venn plots compare DEG mobilized by IL-7 in (F) naïve or (G) memory T cells. (H) Positively regulated DEG from (E) were subjected to hypergeometric testing against the indicated databases (per Fig. 4C). Scatter plot shows enrichment *q* values for top STAT5-regulated pathways (X axis) across genotypes, cytokines and cell states (Y axis; blue = AAB versus WT, orange = BBA versus WT). All gene sets catalogued in Table S1 and Table S5. Gene set usage is detailed in Table S2. Ex vivo culture conditions are detailed in Table S7. Replicate counts and statistical tests for all experiments are listed in Table S6.

**Fig. S6.**
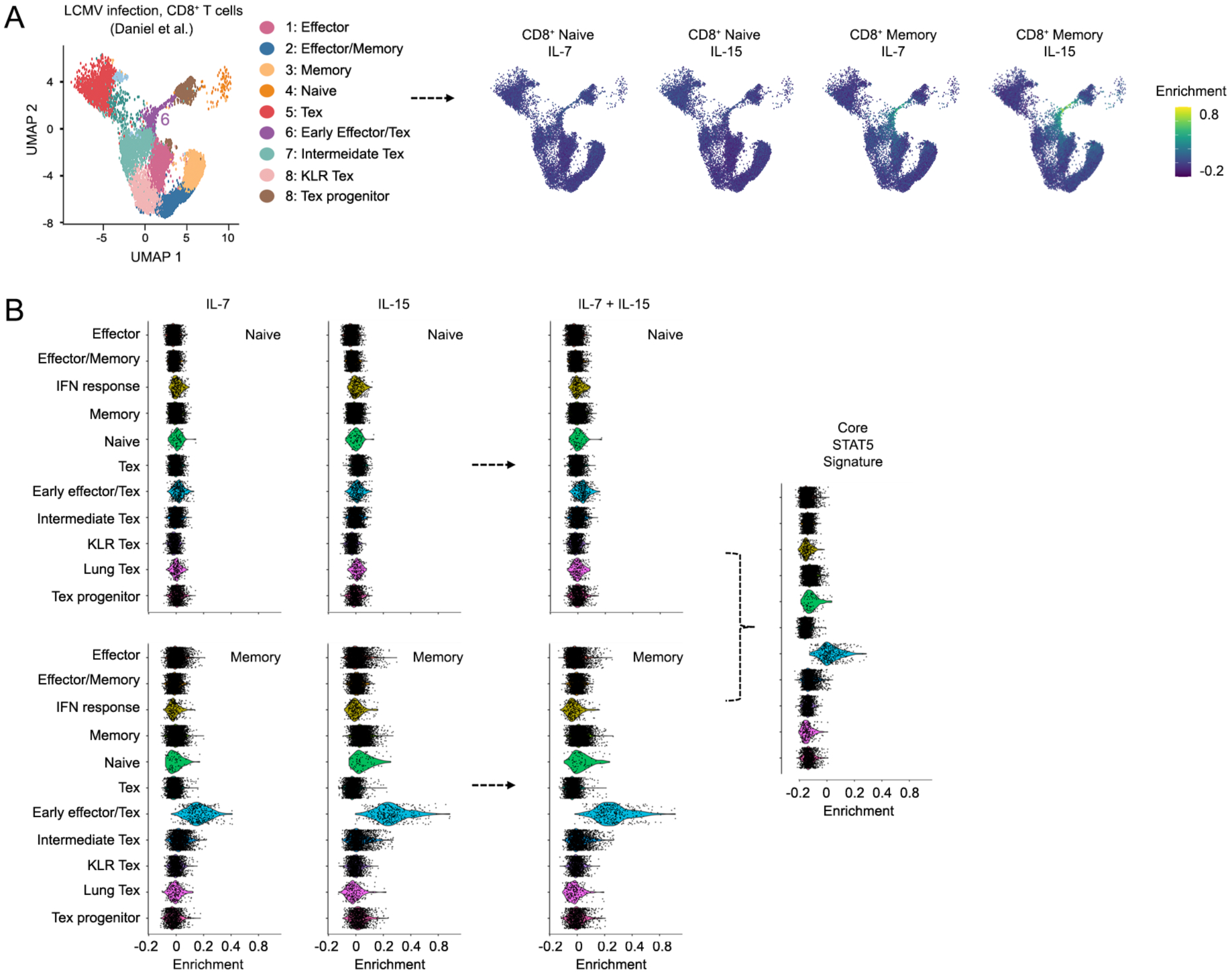
Core STAT5 signature applied to sc-RNAseq data. (A) scRNA-seq UMAP projection segregates CD8^+^ T cells responding to acute or chronic LCMV infection. Clustering and annotation are as published. Feature plots show module score enrichment for the indicated gene signatures. (B) Violin plots show enrichment of STAT5 signatures within individual cells, across clusters. All gene signatures catalogued in Table S5. Replicate counts and statistical tests for all experiments are listed in Table S6.

